# HacDivSel: Two new methods (haplotype-based and outlier-based) for the detection of divergent selection in pairs of populations

**DOI:** 10.1101/026369

**Authors:** A. Carvajal-Rodríguez

**Affiliations:** Departamento de Bioquímica, Genética e Inmunología. Universidad de Vigo, 36310 Vigo, Spain.

## Abstract

The detection of genomic regions involved in local adaptation is an important topic in current population genetics. There are several detection strategies available depending on the kind of genetic and demographic information at hand. A common drawback is the high risk of false positives. In this study we introduce two complementary methods for the detection of divergent selection from populations connected by migration. Both methods have been developed with the aim of being robust to false positives. The first method combines haplotype information with inter-population differentiation (*F*_ST_). Evidence of divergent selection is concluded only when both the haplotype pattern and the *F*_ST_ value support it. The second method is developed for independently segregating markers i.e. there is no haplotype information. In this case, the power to detect selection is attained by developing a new outlier test based on detecting a bimodal distribution. The test computes the *F*_ST_ outliers and then assumes that those of interest would have a different mode. We demonstrate the utility of the two methods through simulations and the analysis of real data. The simulation results showed power ranging from 60-95% in several of the scenarios whilst the false positive rate was controlled below the nominal level. The analysis of real samples consisted of phased data from the HapMap project and unphased data from intertidal marine snail ecotypes. The results illustrate that the proposed methods could be useful for detecting locally adapted polymorphisms. The software HacDivSel implements the methods explained in this manuscript.

## Introduction

Current population genetics has an important focus on the detection of the signature of natural selection at the molecular level. The detection of the selection effect in a given DNA region is important because it may connect that region with a key functionality, or past or ongoing selective events, to have a deeper understanding of the evolutionary processes. Indeed, it may also help to understand the evolutionary mechanisms allowing the species adaptation to local conditions. The study of local adaptation processes implies that some genetic variant is favored by the environmental conditions. The positively selected locus increases in frequency and the pattern of variation around that locus will change, a process known as selective sweep [1, 2].

There are many tests designed for detecting different kinds of effects produced by selective sweeps. Such effects can involve skewed site frequency spectra, high linkage disequilibrium or high rates of genetic divergence [3, 4]. The information required by those tests is variable: they could need knowledge about candidate adaptive loci, their haplotypic phase, their recombination rates, or the ancestral/derived status at each segregating site. This kind of information is often available for model organisms and so, in previous years, most of the effort was focused in humans and other model species.

For non-model organisms, the most useful methods for studying local adaptation have been those based on measuring genetic differentiation between populations. The idea behind these methods is that the loci involved in local adaptation would be outliers, i.e. would have unusually large values of *F*_ST_. From its original formulation [LK test 5], this technique has been improved in several ways depending on the summary statistic’s -the *F*_ST_ or any other differentiation index - expected neutral distribution. That is, in order to account for more realistic situations, the different methods change the assumptions of the null demographic model [reviewed in 6].

However, one of the main drawbacks of the outlier-based methods is the difficulty of defining an accurate null model since there are several historical events and demographic scenarios (other than local selection) that can produce similar *F*_ST_ patterns. Even in the presence of local adaptation, we can expect different *F*_ST_ patterns. This is because the involved populations may be more or less connected by migration, and this event will influence the structure of the genetic variation both at intra and inter-population levels. Consequently, the outlier based methods tested against the deviation over an expected demographic null model, always face the risk of having an excess of false positives [7-10].

Another problem concerning outlier-based methods is that they have low power when the overall *F*_ST_ is high as it can happen when the genetic basis of the adaptation is polygenic (since each particular gene may not have a strong difference with the overall *F*_ST_, hindering the detection of outliers), or when the populations under study are subspecies [6, 11]. Fortunately, over recent years, the amount of information available on the genomes of several species has increased [12] and consequently new and more sophisticated methods can now be applied to detect local adaptation in non-model organisms.

As mentioned above, the linkage disequilibrium (LD) is the basis for several computational methods used in the detection of selective sweeps [reviewed in 13]. Some LD-based methods try to identify maximized LD regions [14, 15] while others explore the pattern of LD decay from different candidate SNPs [16, 17]. However, only a few LD-based methods have considered structured populations as the evolutionary scenario of interest. Still, there are different scenarios that can be evaluated under structured populations [18]. Therefore, although LD-based tests can be powerful and robust for detecting selective sweeps (in isolated or simple structured population scenarios) with low migration rate, they fail to detect it, under several other realistic scenarios [3, 13, 18, 19].

In addition, the possibility of observing local adaptation with gene flow depends on the demography, and on the genetic basis of the traits involved [20]. This decreases the performance of the methods under moderate-to-high migration scenarios which may result into high rate of false positives [18]. Thus, even if haplotype phase information is at hand, specific methods should be developed to detect local adaptation under structured population scenarios [19].

The aim of this paper is to present two complementary methods (for linked and unlinked markers) specialized for detecting divergent selection in pairs of populations with gene flow. The main advantage of both methods proposed here over the existing ones is that they were especially designed to be robust to false positives. Both methods are suitable for working with non-model species, although the first (linked markers) requires an approximate knowledge of the phase of the SNPs under study. Notably, this method still performs quite well under 10-40% of phasing error depending on the marker density. If the SNPs under evaluation are not linked, then the second method should be applied. Our working definition of non-model species includes those for which we could have partial information about the haplotypic phase but a) no estimates of recombination rates, b) no information on potentially adaptive loci and c) knowledge the ancestral/derived status at each segregating site. This definition also implies that we barely know the demography of the populations under study, and if so, we cannot reliably use a simulated neutral distribution to assess significance.

The first method (*nvdF*_*ST*_) combines haplotype-based information with a diversity-based *F*_ST_ measure. It is a sliding-window approach that uses automatic decision-making () to apply different window sizes. On the other hand, the second method (EOS) is not haplotype-based but performs a two-step *F*_ST_ outlier test. The first step of the algorithm consists of a heuristic search for different outlier clusters, while the second step is just a conditional LK test that takes place only if more than one cluster is found. In this latter case, the test is applied through the cluster with the higher *F*_ST_ values.

The design of the work is as follows: in the first part of the article the rationale of the methods is explained. After that, we give the results of the application of both methods to the simulated scenarios. This permits to appreciate their performance in terms of power and number of false positives. Finally, the *nvdF*_*ST*_ method is applied to genome-wide phased data from the HapMap project [21]. This allows to check its performance compared with other methods previously applied to the same data. Additionally, the EOS test is applied to a recently published data set of *Littorina saxatilis* species. More extensive details and a full mathematical description of the methods is given in the Appendix (see S1_Appendix file).

## Models and Methods

### The nvdF_ST_ model

In this section we improve a previous haplotype-based method to detect divergent selection [19, 22]. The new statistic is called *nvdF*_*ST*_ because it combines a normalized variance difference (nvd) with the *F*_ST_ index. The *nvd* part performs a sliding-window approach to identify sites with specific selective patterns. When combined with the *F*_ST_, it allows the significance of the candidate sites to be assessed without the need of simulating neutral demography scenarios. Before developing the *nvd* formula, we review some concepts related to haplotype allelic classes.

### Generalized HAC variance difference

A major-allele-reference haplotype (MARH) is a haplotype that carries only major frequency alleles [22]. Therefore, we can define the mutational distance between any haplotype and MARH as the number of sites (SNPs) carrying a non-major (i.e. minor) allele. Each group of haplotypes having the same mutational distance will constitute a haplotype allelic class (HAC). Therefore (with some abuse of notation) we also call HAC to the value of the mutational distance corresponding to each haplotype allelic class. That is, every haplotype having the same number of minor alleles belongs to the same HAC class and its HAC value corresponds to the number of minor alleles it carries.

Given the definitions above, consider a sample of *n* haplotypes of length *L* SNPs. For each evaluated SNP *i* (*i*ϵ[1,*L*]) we can perform a partition of the haplotypes (and their HAC classes) into *P*_1_, the subset carrying the most frequent (major) allele at the SNP *i* and *P*_2_ the subset with the remaining haplotypes carrying the minor allele at *i*. That is, let ‘0’ to be the major allele for the SNP *i* and ‘1’ the minor. Then, *P*_1_ includes every haplotype carrying the allele ‘0’ for the SNP *i* and *P*_2_ the remaining haplotypes carrying ‘1’ for that SNP. In *P*_1_ we have different HAC values depending on the distance of each haplotype from MARH and similarly in *P*_2_. Within each subset we can compute the variance of the HACs. Thus, in *P*_1_ we have the variance *S*^2^_1i_ and correspondingly variance *S*^2^_2i_ in *P*_2_ where *i* refer to the SNP for which we have performed the partition.

The rationale of the HAC-based methods relies on the sweeping effect of the selectively favored alleles. Therefore, if the SNP *i* is under ongoing selection then the variance in the partition 1 (*S*^2^_1i_) will tend to be zero because the allele at higher frequency (i.e. the allele of the SNP *i* in the partition 1) is being favored and the sweeping effect will make the HAC values in this partition to be lower (because of sweeping of other major frequency alleles) consequently provoking lower variance values [22]. The variance in the second partition (*S*^2^_2i_) should not be affected by the sweeping effect because it does not carry the favored allele. So, the difference *S*^2^_2i_ - *S*^2^_1i_ would be highly positive in the presence of selection and not so otherwise. For a window size of *L* SNPs, the variance difference between *P*_2_ and *P*_1_ can be computed to obtain a summary statistic called Svd [22] that can be generalized to

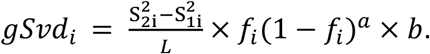
Where *f*_i_ is the frequency of the derived allele of the SNP *i*, and the parameters *b* and *a* permit to give different weights depending on if we need to detect higher frequencies (*a* = 0) or more intermediate ones (*a* > 0) of the derived allele. If *a* = 0 and *b* = 1 the statistic corresponds to the original Svd and if *a* =1 and *b* = 4 it corresponds to the variant called SvdM [19]. Note that when using *a* = 1, it is not necessary to distinguish between ancestral and derived alleles because *f*_i_ and 1-*f*_i_ are interchangeable.

A drawback in the gSvd statistic is its dependence on the window size as it has already been reported for the original Svd [19, 22]. Although gSvd is normalized by *L*, the effect of the window size on the computation of variances is quadratic (see Appendix A-1) which explains why the normalization is not effective in avoiding a systematic increase of the statistic under larger window sizes. This bias, due to the change in the window size, is important because the partitions *P*_1_ and *P*_2_ may experience different scaling effects, which would increase the noise in the estimation. The change in the scale due to the window size will also be dependent on the recombination and selection rates. Thus, it is desirable to develop a HAC-based statistic that does not increase with the window size. Following, the between-partition variance difference is reworked in order to develop a new normalized HAC-based statistic, specially focused on detecting divergent selection in local adaptation scenarios with migration.

### Normalized variance difference (nvd)

We have seen that in a sample of *n* haplotypes, we can compute the statistic gSvd which basically is the difference between the HAC variances of partitions *P*_1_ and *P*_2_. The corresponding HAC means and variances at each partition are related via the general mean and variance in that sample. Considering, for any candidate SNP, *m* the mean HAC distance of the sample, and *m*_1_ and *m*_2_ the means of the partitions *P*_1_ and *P*_2_, respectively. We have the following relationships for the mean *m* and the sample variance *S*^2^ (the subscripts 1 and 2 identify their corresponding partitions *P*_1_ and *P*_2_; see the Appendix A-2 for details)

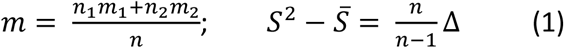
With 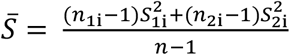; *n*_1_ and *n*_2_ are the sample sizes at each partition and 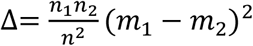.

Using the relationships in (1), it is possible to compute the variance difference as appears in gSvd. However, we can substitute the parameters *b* and *a* by *a* = 1 and *b* = 4 as these are the values that permit to ignore the allelic state while maximizing the frequency product. It is also possible to consider the difference between means term (delta) in order to engage it in the detection of selection (see details in the Appendix). Thus, we finally obtain a new statistic for the variance difference of the candidate SNP *i*

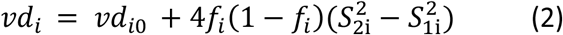
where 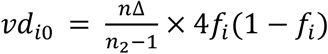 The effect of selection upon *vd*_i_ is two-fold. The first term of the sum in (2) corresponds to the effect upon the difference between means and the second term between variances. Increasing *S*_2i_ or decreasing *S*_1i_, as expected under selection, will increase the value of the statistic. If *S*_1i_ and *S*_2i_ are equal, then the value of *vd*_i_ is independent of the variances and only relies on the term *vd*_i0_ corresponding to the partitions’ mean (*m*_1_ and *m*_2_) and the candidate SNP frequencies.

It is worth mentioning that the two parts of *vd*_i_ are not independent. If we have an extreme value for the HAC mean in the selective partition, for example *m*_1_ = 0, this implies that *S*_1_^2^ = 0 since every haplotype has to have a HAC of 0 to get that mean value. Note however, that the opposite is not true: a value of *S*_1_^2^ = 0 does not imply necessarily that *m*_1_ = 0.

At intermediate frequencies, an upper bound of (2) is (see Appendix A-2 for further details)

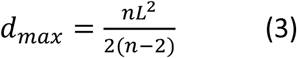
If we divide (2) by *d*_max_, we have a normalized variance difference

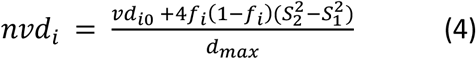
In a two-populations setting, it is possible to combine the sequences from the two populations in a unique sample and so the quantity from (4) can be computed for each SNP in that sample. The SNP yielding the maximum *nvd* may be considered as a candidate for divergent selection. When pooling both populations, the SNP frequencies tend to be intermediate in the divergent selective sites.

Therefore, the calculation of *nvd* consists of merging the shared SNPs from the two population samples, and then computing the normalized variance difference using (4). Because selection acts at the intrapopulation level the reference haplotype (MARH) is defined just from one of the populations. The *nvd* neutral distribution so calculated under different window sizes can be viewed in the supplementary Figure S1, Appendix A-2. Under low window size, *nvd* has higher variance and is slightly biased toward positive values. The effect of increasing the window size is a slight reduction of the *nvd* mean value and variance.

Provided that the sample size is the same in both populations, the choice of the reference does not have an appreciable effect either in the power or in the false positive rate of the test (supplementary Figure S2 and Table S1). However, if the sample sizes are different, then the reference should come from the population with the highest sample size; if not, the method can suffer an important loss of power.

We have still pending the problem of choosing an optimal window size. A possible solution is to automate its choice by selecting the size which gives the maximum value for the statistic [19], or alternatively, by trying different window sizes and giving the corresponding results for each window at the output. We have opted for the latter and so, under a given window size, the accompanying program evaluates all the SNPs and selects the ones that have the maximum *nvd* (one or more depending on the proportion of candidates we are interested in evaluating) and then, repeats the process for a different window size.

At this point we already have a HAC-based statistic, *nvd*, that does not increase with the window size and should produce higher positive values for pairs of populations undergoing divergent selection (see Figure S2). However, even if there is no selection, the maximum *nvd* value could also be positive (see Figure S1).

Unfortunately, the neutral distribution still depends on the combined effect of window size, marker density, and recombination. Besides, we ignore the theoretical distribution of the statistic and cannot decide if a given maximum is supporting the hypothesis of selection or not.

In addition, we might not have enough information on the species to simulate its evolution under a given neutral demography. Therefore, we still need to identify whether the value obtained for a given sample is due to the effect of selection, particularly because we want to put a great emphasis on avoiding false positives.

Consequently, we incorporate two more measures before giving a diagnosis about the presence of divergent selection. The first is a sign test based on the lower bound of nvd, the second is the comparison between the *F*_ST_ of the SNP having the maximum *nvd* and the overall *F*_ST_.

### Sign test

In our definition of *nvd* the term *vd*_i0_ cannot be negative. For that reason, under some scenarios and window sizes, *nvd* is biased toward positive values. In such cases, a negative (or null) variance difference could be linked to a high positive *vd*_i0_ term provoking a positive *nvd* in a clearly neutral case. Therefore, we use a lower bound of *nvd* to derive the quantity called selection sign, *ssig* (see Appendix A-3) that would have negative values when the HAC values in the first partition are high which is not expected under selection

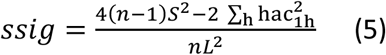
where hac_1h_ are the HAC values measured at each haplotype *h* in the partition 1 and the sum is over the *n*_1_ sequences in that partition. A negative sign in (5) suggests that the value of *nvd* is not the result of divergent selection (see supplementary Figure S2). Indeed, we require (5) to be positive to count a given candidate as significant.

Finally, even if we have a candidate position identified by its high *nvd* value and by the positive sign in *ssig*, we still lack a method for obtaining p-values associated to the sites chosen by the *nvd* algorithm. We can solve this problem by combining the information on the selective candidate SNP, as given by nvd, with the *F*_*ST*_ interpopulation differentiation index at that site. The joint use of these methods produces the combined measure *nvdF*_*ST*_.

### Combined method: *nvdF*_ST_

First, it is important to note that when computing *nvd*, we considered only the SNPs shared between both populations in order to avoid low informative loci with high sampling variance [reviewed in 6]. Thus, we have an *nvd* value that may indicate the presence of divergent selection in a pair of populations connected by migration. The rationale of the *nvdF*_ST_ approach is that if divergent selection acts on a specific site then the *F*_ST_ at that site would be higher when compared to the overall *F*_ST_. Therefore, we proceed as follows, let *i* be the candidate site chosen because it has the maximum *nvd* value, then we calculate the index *I*_i_ = *F*_STi_ − *F*_ST_ that compares the *F*_*ST*_ measured at that site with the overall. The *F*_ST_ values were computed following the algorithm in Ferretti *et al* [23]. To obtain the p-value, we do not perform an LK test [5] because first, the candidate was not chosen for being an outlier and second, we are considering linked rather than independent sites.

To get the *p*-value for a given index *I*_i_, the data is resampled several times (500 by default, and 100 when we evaluate more than 10^4^ SNPs) to generate an empirical distribution. The expected frequency of each SNP is obtained as its mean frequency between populations, as this is the expectation under the homogenizing effect of migration [24]. If the sample sizes are different, the mean is weighted by the sample size. Then, for each resampling iteration, the probability of a given allele at each population is obtained from a binomial *B*(*p*,*n*), where *p* is the mean allelic frequency at that site, and *n* the local population sample size. The p-values correspond to the proportion of times that the resampled indexes (*I*^’^_i_ = *F*_STi_resample iteration_ − *F*_ST_resample iteration_) were larger than the original index *I*_i_.

We have computed the p-values for all the SNPs. If we inspect all of them and ignore the sign test, the above procedure is just testing the hypothesis of panmixia for each SNP. However, we are not doing so, i.e. we are not choosing the most significant *p*-values and so we are not targeting SNPs that reject panmixia. On the contrary, we are selecting some candidate SNPs based on their high *nvd* value and positive sign test. Only then, we check if such SNPs reject panmixia; if so, this adds evidence for divergent selection since it is well-known that divergent selection rejects that hypothesis for the selective genes [25, 26]. Regarding the number of candidates ranked by their high *nvd* value, we can decide to consider just the best one, a few, or a given percentage (say 0.1%) of candidates.

For candidates with similar frequencies at both populations, we expect low values for the index *I*_i_ and correspondingly high *p*-values. When their pooled frequency is intermediate; two situations are possible: first, each population has a similar intermediate frequency which again implies high *p*-values; or second, the frequencies can be extreme and opposite at each population. In the latter, *I*_i_ is high and its *p*-value low. Note that, for each site, the resampling procedure has a variance *p*(*1*-*p*)*n* which is large at intermediate pooled frequency values.

Thus, this method has the desired property to be more conservative at intermediate pooled frequencies, which minimizes the possibility of false positives.

### Significance, effective number of independent SNPs and *q*-value estimation

The computation of *nvdF*_*ST*_ required as many tests as the candidates considered under the different window sizes (*N*_w_) assayed. Thus, it is appropriate to apply a multiple testing correction for the number of candidates (*N*_*C*_).

The Šidák correction [27, 28] can be applied to get the adjusted significance level s_L_ = 1 − (1 - *γ*)^1/*C*^ with nominal *ϒ* (= 0.05 by default). The value of *C* is *C* = *max*{(θ_πmax_ − θ_π_)/(θ_πmax_ − θ_πmin_)×*L*_min_, (*N_C_* + *N*_w_)} where θ_π_ is the average number of differences between pairs of sequences computed for a given window size. If sample size *n* is even then θ_πmax_ = n/(2*(n-1)) and θ_πmin_ = 2\**maf*\*(*n*-*maf*) / [*n*\*(*n*-1)] *maf*. The rationale for that *C* value is that in the case of low diversity samples it could happen that there are only one window size above the minimum (*L*_*min*_). Low window sizes are precisely those where the *nvd* statistic is more biased toward positive values (Figure S1). To avoid false positives in the case of only one candidate and low diversity, we correct by the minimum window size, as this involves the minimum number of SNPs tested but weighted them by (θ_πmax_ − θ_π_)/(θ_πmax_ − θ_πmin_) which tends to 1 when diversity is very low or, 0 if high.

Thus, the algorithm *nvdF*_*ST*_ would finally suggest a candidate as significant only when the sign, as computed in (5), was positive, and the p-value (as obtained in the previous section) is lower than *s*_L_.

In addition, it could be of interest to have information about the number of independent SNPs included between the left and right-most candidate positions. To roughly estimate the number of independent SNPs, we calculated the linkage disequilibrium measure *D*’ [29, 30] at each pair of consecutive sites, and then store the quantity *r*’ = 1 − |D’| for each pair. The effective number of independent SNPs (*M*_effs_) between site *w*_ini_ and *w*_end_ is then obtained as one plus the summation of the *r*’ values in the interval [*w*_ini_, *w*_end_).

The *q*-values [31] can be seen as multiple testing analogs of *p*-values. They have been proposed as a useful approach for evaluating method performance in terms of false discoveries [11]. Accordingly, we estimate the *q*-values (see reference [32] and Appendix A-4 for details on the calculation) and provide those corresponding to each inspected *p*-value. In order to obtain the *q*-values we must compute the *p*-values for all the SNPs (shared SNPs at frequency higher than *maf*). Thus, all the *p*−values and the corresponding *q*−values are calculated. Only those corresponding to the desired number of candidates, ranked by their highest *nvd* values, are given. For example, consider that the highest *p*-value is 0.99 and the lowest is 10^-6^. If we select just one candidate is because it has the highest *nvd* value and we concern only a posteriori about its *p*-value. If the associated *p*-value happens to be 0.2, we give both the *p*-value and its corresponding *q*-value, producing an output of this as a non-significant result.

### The extreme outlier set test (*EOS*)

The *nvd*F*_ST_* method assumes the existence of linked genetic markers. If the data consists mostly of independent markers this would provoke a failure to detect the selection pattern because the HAC-based information does not exist. To deal with this situation, a second method was implemented consisting of a two-step heuristic procedure that performs a conservative test for identifying extreme outliers.

As already mentioned, the variance of the *F*_ST_ distribution is quite unpredictable under a variety of scenarios. This provokes high rates of false positives associated with the *F*_ST_ outlier tests. Our heuristic strategy takes advantage that independently of the demographic scenario, the involved regions under divergent selection may produce extreme outliers that would be clustered apart from the neutral ones. The subsequent LK test is performed only when this kind of outliers is detected. As *F*_ST_ estimator we use *G_ST_* [33].

The rationale of the algorithm is as follows: the first step consists of computing the extreme positive outliers in the sense of Tukey, i.e. those sites having a *F*_ST_ value higher than 3 times the interquartile range [34]. The second step identifies different classes inside the extreme outlier set. This is done by a *k*-means algorithm [35, 36]. The algorithm permits to classify all the elements of the outlier set in one of the *k* classes. Once all the elements are classified, the class with lower values is discarded. Only the elements, if any, in the upper classes having values higher than a cutoff point are maintained in the set. For the sake of computational efficiency, we use *k* = 2 and consider two modes {0, *F*_STu_} for computing the cutoff. The modes corresponding to lower (0) and upper (*F*_STu_) bounds for the *F*_ST_ estimator (see Appendix A-5). The cutoff point is defined as the overall *F*_ST_ + σ*F*_STu_ / 3, i.e. the mean plus sigma times the square root of the upper-bound for the *F*_ST_ variance under an asymmetric unimodal distribution [37]. The value of sigma was set by default to σ =1.25. Finally, for each of the candidates remaining in the EOS after the cutoff, the LK test [5] is performed to compute its *p*-value. The Šidák correction [27, 28] for the number of remaining outliers in the set is applied to get the significance level. Each *p*-value is accompanied by its corresponding *q*-value (computed as explained in the previous section).

### Software description

Both *nvd*F*_ST_* and the EOS test have been implemented in the program HacDivSel. Complete details of the software and extensive explanations for its use can be found in the accompanying manual. The input program files for the haplotype-based test can be in the SNPs × haplotypes HapMap3, MS [38] or Fasta formats. If the data does not include haplotype information, then the Plink [39] flat file (map/ped), Genepop [40] or BayeScan [41] formats can be used. When using the Fasta format, the sample size should be the same for both populations. A typical command line for calling the program in order to analyze a MS format file named *sel.txt* containing 100 sequences, 50 from each population, would be

> *HacDivSel-input sel.txt -sample 50 -sample2 50 -candidates 10 -SL 0.05 -output anyname -format ms –maf 4*

Where the label, “-*candidates* 10”, indicates that the ten highest *nvd* values should be included in the output. The program analyze the file and produce as output the highest 10 values and their significance at the 0.05 level for different window sizes. It also performs the EOS test and gives the resulting outliers, if any, and their significance. Only the SNPs shared by the two populations are considered. This implies that there are at least 4 copies of each SNP in the metapopulation (maf = 4). The set of command-line options utilized for the different analyses, both for simulated and real data in the next sections is available in the S2_Sim file.

### Simulations

There are several examples of adaptation to divergent environments connected by migration, e.g. the intertidal marine snail *L. saxattilis* [42], some wild populations of *Salmo salar* [43], Coregunus species [44], etc. To perform simulations as realistic as possible, we use some relevant demographic information from *L. saxatilis*, such as migration rates and population size as estimated from field data [42]. Concerning selection intensities, we considered moderate selection pressures and few loci with large effects [45]. Therefore, a model resembling the most favorable conditions for the formation of ecotypes under local adaptation with migration was implemented.

Two populations of 1000 facultative hermaphrodites were simulated. The selective scenario (*α* = 4*Ns*) is divergent so that the allele favored in one population is the deleterious in the other. Each individual consisted of a diploid chromosome of length 1Mb. The contribution of each selective locus to the fitness was 1-*hs* with *h* = 0.5 in the heterozygote or *h* = 1 otherwise. The selection coefficient for the ancestral allele was always *s* = 0 while *s* = ± 0.15 for the derived. That is, the ancestral was the favored allele in one population (positive *s* in the derived) while the derived was the favored in the other population (negative *s*, see supplementary Table S2 in Appendix A-6). In the polygenic case the fitness was obtained by multiplying the contribution at each locus. This simulation model involves low and high mutation rate (θ=4Nμ ϵ {12, 60}) and different recombination rates (ρ = 4*Nr* ϵ {0, 4, 12, 60, unlinked}) and extends previous work [19] by adding new parameter values and demographic scenarios. At the end of each run, 50 haplotypes were sampled from each population. The whole setting is fully explained in the Appendix (Appendix A-6). The simulations were performed using the last version of the program GenomePop2 [46].

## Results

We present here the results for both simulated and real data. The power of a test (true positive rate, TPR) is measured as the percentage of runs in which selection was detected from simulated selective scenarios, and the false positive rate (FPR) is measured as the percentage of runs in which selection (at any position) was detected from simulated neutral scenarios. The *q*-value [31] is estimated from the results (see Appendix A-4).

The simulated data is also utilized for comparing *nvdFST* with the related statistic SvdM (that needs a simulation step to assess significance) while the EOS test is compared with BayeScan 2.1 [41]. We chose BayeScan because it is one of the main state-of-the-art outlier-based programs. The parameters for BayeScan were the default ones as this is a conservative setting and we were interested in comparing the false positive rates. Only SNPs shared between populations and with a minimum allele frequency (maf) of 2 per population (4%) were considered.

### Precision

The precision or positive predictive value (PPV) is a useful measure for answering the question of how well a positive result in the test predicts the real existence of selection. It is calculated as the proportion of true positives (TPR) out of all positive results (TPR + FPR).

Recall that one of the main focus of the proposed methodologies is to have low false positive rate (FPR) and, from this point of view, the PPV measure is very informative since the higher the PPV, the lower the possibility that a positive result be a false one.

However, if a test has very low false positives, it could have high PPV but low power. In this case, the high PPV is hiding the low performance of the test. Thus, we are interested in both, the PPV as shown in the Figure 1, but also in the specific values in terms of power (TPR) and false positives (FPR), as given in the text and in the tables following.

Concerning the PPV measure, the rate of true positives with respect to the total positives is above 90% for the haplotype based methods *nvd*F*_ST_* and *SvdM* [19], as shown in Figure 1 (left panel). The simulated scenarios were different combinations of mutation and recombination rates using *α* = 600 and *Nm*=10 (Appendix A-6).

The individual TPR and FPR values for *nvd*F*_ST_* are given in Table 1. For *SvdM*, the TPR had a minimum of 43% and a maximum of 94%. The FPR is fixed to 5% since this was the critical threshold used by the authors in the neutral simulations for assessing significance. The values used for computing the *SvdM* points in the Figure correspond to those in Tables 2 and 3 in [19]. The general performance of *SvdM* was slightly worse than for *nvd*F*_ST_*.

**Table 1.** Performance of the combined method (*nvd*F*_ST_*) with *n* = 1 selective site located at the center of the chromosome or *n* = 5 (see Appendix A-6). Selection was *α* = 4*Ns* = 600 and migration *Nm* = 10. Mean localization is given in distance (kb) from the real selective position.

| $\Sigma$ | $\theta$ | $\rho$ | $n$ | %Power | %FPR ($\gamma = 5\%$) | $q$-value | Localization (kb) |
| --- | --- | --- | --- | --- | --- | --- | --- |
| 65 | 12 | 0 | 1 | 87 | 2.1 | 0.0058 | $\pm 458$ |
| 63 | 12 | 4 | 1 | 94 | 2.7 | 0.0008 | $\pm 200$ |
| 60 | 12 | 12 | 1 | 90 | 1.0 | 0.0003 | $\pm 33$ |
| 251 | 60 | 0 | 1 | 79 | 1.8 | 0.0048 | $\pm 60$ |
| 232 | 60 | 4 | 1 | 84 | 6.2 | 0.0011 | $\pm 17$ |
| 249 | 60 | 60 | 1 | 86 | 2.4 | 0.0002 | $< \pm 1$ |
| 282 | 60 | 60 | 5 | 99 | 2.4 | 0.0002 | $< \pm 1$ |
| 318 | 60 | ∞ | 1 | 0 | 0 | - | - |

$\Sigma$ : Mean number of shared SNPs per Mb. $\theta$ : Mutation rate. $\rho$ : Recombination rate. FPR: false positive rate. $q$ -value: mean estimated $q$ -value for the significant tests. ∞ : Independently segregating sites.

The non haplotype-based methods, EOS and BayeScan (BS), have no power when the markers are linked, so they are compared only under independent and weak-linkage (ρ = 60; 1.5 cM/Mb) scenarios (right panel in Figure 1). As it can be seen in Figure 1, the EOS method has the best performance in terms of the proportion of true positive results from the total positives.

For EOS, the TPR was about 60% with a very low FPR that provokes that almost 100% of positives are true ones. The TPR for BS with Bayes factor of 3 (BSBF3) was 47% under weak linkage and 93% with independent markers. However, both settings had high rate of false positives with FPR of 26 and 57% respectively. The TPR of BS with Bayes factor of 100 was only 13% (FPR 3%) under weak linkage but was 82% (FPR 8%) under independent markers.

**Fig 1.** Positive predictive values for *nvdF*_ST_, SvdM, EOS and BayeScan (Bayes factor 3: BSBF3 and Bayes factor 100: BSBF100) methods. There are 6 points (2 mutation x 3 recombination) in the curves corresponding to the haplotype-based method (left panel) and 2 points (high recombination and independent markers) in the outlier-based method (EOS and BayeScan, right panel).

The above results are observations from 1,000 runs. Within each run, we have also measured the number of sites falsely detected per genome, as well as the FDR (the proportion of falsely detected sites from the total detected). As these quantities were low and do not have any significant impact on power (TPR) and FPR (as defined above), we will postpone the mention of the within-run measures until the discussion section.

In the following sections, we detail the results for *nvd*F*_ST_* and EOS under different simulated scenarios.

### Combined Method (*nvdF*_ST_)

Under a single locus architecture with selection *α* = 4*Ns* = 600 and migration *Nm*=10, the power of *nvd*F*_ST_* vary between 79-94% for both medium (60 SNPs/Mb) and high density (250 SNPs/Mb) maps (Table 1). These results can be compared to published analyses [19] with the methods Svd and SvdM for which slightly worse performance (42-94%) were obtained for the same mutation and recombination [Tables 2 and 3 in 19]. However, as mentioned elsewhere, the methods Svd and SvdM oblige the user to perform simulations of the neutral demography to obtain the *p*-values for the tests, and consequently, the results in the Rivas and coworkers study [19], were obtained having the exact neutral demography available. As it can be appreciated from rows 1 to 6 in Table 1 —that matches the scenarios in [19]— the *nvd*F*_ST_* performs well without the need of performing additional neutral simulations. Also, the false positive rate and the *q*-value (mean *q*-value through significant runs) are low in all the scenarios. The shown results are for 10,000 generations; the cases with 5,000 generations were similar.

Under the polygenic architecture (*n* = 5 in Table 1) at least one candidate is found 99% of the time, and more than one, 80% of the time. However, the number of correctly identified sites is quite variable ranging between 1 and 3.

The last row in Table 1 corresponds to the case when all SNPs segregate independently. In this case, the method fails to detect selection. This is not surprising because the information from the haplotype allelic classes is absent under linkage equilibrium; the adequate patterns are not found, which provokes both a negative in the sign test and a candidate with low *F*_ST_ index measure.

### Phasing accuracy

We have tested the impact of phasing error by introducing different percentage of random error in the allele imputation. At each sequence, 10-50% of imputation errors were introduced in random positions. This process was performed for the selective cases *α* = 600 with θ = 60 and ρ = {0, 4, 60} corresponding to the same cases (with 100% accuracy) from rows 4-6 in Table 1. The obtained data were analyzed with *nvd*F*_ST_* under different window sizes (automatic mode). The best performing window size was *L* ϵ {55,75}.

**Fig 2.** Effect of % phasing accuracy on the power of the *nvd*F*_ST_* test.

From Figure 2 it seems that *nvd*F*_ST_* is robust up to 10% of imputation error (90% accuracy). The explanation is that a given error rate will affect more at specific window sizes (higher window sizes), because the automatic window size selection method tries different lengths so that the shortest window maintain the power. Coherently with this explanation the more affected cases correspond to a higher recombination rate and window size.

Still, when the recombination rate is not high, *nvd*F*_ST_* seems to be robust under imputation error as high as 20 to 30%. Finally, a phasing error of 50% provokes the method failure, independently of the linkage relationship. Thus, under imputation error of 10-30%, the more dense the map of markers is, the more robust the method. Hence, under high-medium density maps, as those we have assayed, the performance is also reliable when using different subsets of SNPs provided that the linkage relationships are not completely broken (see supplementary Figure S3).

### Short-term Strong Selection and Long-term Weak Selection Scenarios

The performance of *nvd*F*_ST_* under the strong selection scenario (*α* = 6000) in the short-term (500 generations) varies between 44%, for fully linked, to 67%, for weak linked markers (Table 2). Not surprisingly, the number of segregating sites is considerably reduced. In fact the minimum window size allowed by the program had to be shortened from 51 to 25 to perform the analyses. Notably the false positive rate (FPR) was 0.

Concerning weak selection in long-term scenarios (Table 2, *α* = 140), the power varies between 36-17% with false positive rates between 1 and 2%.

**Table 2.** Performance of the combined method (*nvd*F*_ST_*) with a single selective site in the short-term strong (*α* = 6000) and the long-term weak (*α* = 140) selection scenarios. *Nm* was 10. Mean localization is given in distance (kb) from the real selective position.

| $\Sigma$ | $\theta$ | $\rho$ | $\alpha$ | $t$ | %Power | %FPR ($\gamma = 5\%$) | $q$-value | Localization<br>(kb) |
| --- | --- | --- | --- | --- | --- | --- | --- | --- |
| 112 | 60 | 0 | 6000 | 500 | 44 | 0 | 0 | $\pm 66$ |
| 32 | 60 | 4 | 6000 | 500 | 63 | 0 | 0.0014 | ±5 |
| 62 | 60 | 60 | 6000 | 500 | 67 | 0 | 0.0008 | ±93 |
| 165 | 60 | 0 | 140 | 5,000 | 36 | 1 | 0.007 | ±2 |
| 156 | 60 | 4 | 140 | 5,000 | 30 | 1.5 | 0.003 | ±13 |
| 135 | 60 | 60 | 140 | 5,000 | 17 | 2 | 0.000 | ±1 |

$\Sigma$ : mean number of shared SNPs per Mb. $\theta$ : Mutation rate. $\rho$ : recombination rate. $t$ : number of generations. FPR: false positive rate. $q$ -value: mean estimated $q$ -value, the mean is computed only through the significant tests.

### Extreme Outlier Set Test (EOS)

We applied the EOS test under the single locus architecture with selection *α* = 600 and migration *Nm*=10. As desired, the test is very conservative with false positive rate below the nominal 0.05 in every case (Table 3). Not surprisingly for an outlier-based method, the test has no power if markers are strongly linked (ρ from 0 to 12) or under a polygenic setting (row with *n* = 5 in Table 3). However, in the cases of independent SNPs, and also with recombination of 1.5 cM/Mb in maps with 250-300 SNPs/Mb, the power rises up to 60%. Therefore, the EOS test is complementary to *nvd*F*_ST_* in the sense that EOS has the maximum power when *nvd*F*_ST_* has the minimum one. This behavior is expected because *nvd*F*_ST_* requires some linkage among markers, while EOS requires the contrary.

**Table 3.** Performance of the extreme outlier test (EOS) with *n* = 1 selective site located at the center of the chromosome or *n* = 5. Selection was *α* = 600 and *Nm* = 10. Mean localization is given in distance (kb) from the real selective position.

| $\Sigma$ | $\theta$ | $\rho$ | $n$ | %Power EOS | %FPR ($\gamma = 5\%$) | q'-value | Localization (kb) |
| --- | --- | --- | --- | --- | --- | --- | --- |
| 65 | 12 | 0 | 1 | 0 | 0 | - | - |
| 63 | 12 | 4 | 1 | 0.2 | 0 | 0.46 | $\pm 3$ |
| 60 | 12 | 12 | 1 | 1.1 | 0 | 0.45 | $\pm 77$ |
| 251 | 60 | 0 | 1 | 0.7 | 0 | 0.10 | 0 |
| 232 | 60 | 4 | 1 | 1.3 | 0 | 0.20 | $\pm 150$ |
| 249 | 60 | 60 | 1 | 58 | 0.4 | 0.5 | $< \pm 1$ |
| 282 | 60 | 60 | 5 | 1.6 | 0.4 | 0.49 | $\pm 5$ |
| 318 | 60 | ∞ | 1 | 61 | 1.2 | $3 \times 10^{-6}$ | $\pm 7$ |

$\Sigma$ : mean number of shared SNPs per Mb. $\theta$ : mutation rate. $\rho$ : recombination rate. FPR: false positive rate. q'-value: mean corrected (see appendix A-4) estimated $q$ -value in the significant tests. ∞ : independently segregating sites.

Concerning false positives, note —in the last three rows of Table 3— the low false positive rates (FPR) that indicate the low percentage of outliers detected as selective in the corresponding neutral scenario. Indeed, in the scarce runs where false positives occur their number was very low (1 false outlier in 1.2% of the runs for independent markers and 1 to 4 false outliers in 0.4% of the runs for linked markers). Thus, the test worked correctly by avoiding false selective sites under the neutral setting.

However, the *q*-value estimates varied a lot depending on the linkage between markers. It can be appreciated that the *q*-value is very low (3×10^-6^) for independently segregating sites, but rise up to 0.5 for the same scenario when markers are linked. The reason behind this lies in the way the *q*-values are computed: the algorithm assumes that a number of independent *S* tests were performed and corrects for *S* tests under this assumption (see Appendix A-4). However, in the case of linked markers, the number of independent tests is smaller and this generates inflated *q*-values.

### Position Effect

The ability to locate the physical position of the selective site increases with the marker density and the recombination rate (Table 1). The given localizations are away (in kilobases) from the correct position and averaged through the runs. Standard errors are omitted since they were low (in the order of hundreds of bases or below 5 kb in the worst scenario of fully linked markers). The *nvd*F*_ST_* method performs well when a) the target site is located at the center of the studied region, b) the selection is not too strong (*α* ≤ 600) and c) the overall recombination rate is at least 0.3 cM/Mb (ρ ≥ 12). In this case, the selective location is estimated, at worst, within 33 kb of distance from the true location (Table 1). In the case with strong selection, the localization is still wrongly assigned under high recombination (Table 2, *α* = 6000). However, this could be due to the low number of segregating sites (only 62 in Table 2).

In addition, it should be noted that the proportion of detected sites that lies at, or close to, the true selective position depends on the linkage relationships among the markers, which is obviously influenced by the recombination rate. Thus, if we consider the distance in centimorgans (cM), our results are not so different between the distinct recombination rates. This means that all detected sites lay within 1 map unit from the true selective position. For example, if we require that any detected SNP should be within 0.15 map units from the true position, then under a recombination rate of 0.3 cM/Mb we accept any detected site that is no more than 500 Kb away from the correct position. The same requirement in the cases under recombination rate of 1.5 cM/Mb implies that we only accept sites that are closer than 100 Kb.

Importantly, the localization is also dependent on where the selective site is placed within the chromosome. The farther from the center, the worse the ability to correctly localize the selective positions (Table 4). In this case, with recombination of 1.5 cM/Mb, the inferred location changes from an almost perfect localization (<1 kb from Table 1) to distances of 10-122 kb, as the target is shifted away from the center. This issue has already been shown for other HAC-based methods [19].

**Table 4.** Performance of *nvd*F*_ST_* and EOS with a single selective site located at different positions. Selection was *α* = 600 and *Nm* = 10. Mean localization is given in distance (kb) from the real selective position. FPRs are the same as in Table 1. *q*-value refers to the mean *q*-value for the significant *nvdF*ST tests.

| $\Sigma$ | $\theta$ | $\rho$ | %Power<br>$nvdF_{ST}$, EOS | Position (kb) | $nvdF_{ST}$ $q$-value | Localization (kb)<br>$nvdF_{ST}$, EOS |
| --- | --- | --- | --- | --- | --- | --- |
| 259 | 60 | 0 | 81, 1 | 0 | 0.0044 | +483, +457 |
| 255 | 60 | 0 | 81, 1.5 | 10 | 0.0049 | +433, +496 |
| 256 | 60 | 0 | 82, 0.9 | 100 | 0.0041 | +350, +413 |
| 255 | 60 | 0 | 78, 0.6 | 250 | 0.0039 | $\pm 194$, $\pm 185$ |
| 230 | 60 | 4 | 75, 2.5 | 0 | 0.0014 | +324, +127 |
| 226 | 60 | 4 | 77, 3.5 | 10 | 0.0016 | +326, +142 |
| 233 | 60 | 4 | 80, 1.8 | 100 | 0.0017 | +227, +140 |
| 229 | 60 | 4 | 83, 1.6 | 250 | 0.0009 | $\pm 123$, $\pm 20$ |
| 262 | 60 | 60 | 63, 93 | 0 | 0.0014 | +122, +40 |
| 261 | 60 | 60 | 68, 91 | 10 | 0.0014 | +113, +34 |
| 257 | 60 | 60 | 81, 84 | 100 | 0.0006 | $\pm 44$, $\pm 6$ |
| 252 | 60 | 60 | 87, 67 | 250 | 0.0004 | $\pm 10$, $\pm 0.06$ |

$\Sigma$ : Mean number of shared SNPs per Mb. $\theta$ : Mutation rate. $\rho$ : Recombination rate. Position: real position of the selective site.

At high recombination rates (1.5 cM/Mb), this problem is partially solved by using the EOS test (Table 4). In this case, the test has high power (67-93%) and localizes well the selective position. In fact, the position of the selective site is almost perfectly estimated (few bases or kb) when the true position is not at the extremes. Even if the target sites are at the extremes, the localization is within 40 kb (see cases with ρ = 60 in Table 4).

In the case of independent markers with the selective site located at the center (Table 3) the localization by EOS was good.

## Other demographic scenarios

### Bottleneck-expansion Scenarios

Bottleneck-expansion scenarios are known to leave signatures that mimic the effect of positive selection. We tested the robustness of the tests by looking for false positives when applying the methods to a bottleneck and expansion scenario under a neutral setting. The bottleneck was simulated by a reduction of one of the populations to 1% of the original size (*N* from 1000 to 10). Afterwards, the population expansion was implemented by increasing the population size following a logistic growth model (see Appendix A-6). The methods performed well: the false positive rate is maintained below the nominal level with 4.6% and 1% for *nvd*F*_ST_* and EOS tests, respectively.

### High Migration Scenario

Scenarios with high migration rate are intrinsically difficult for detection of selection. Under the short-term (500 generations) scenario with *Nm* = 50 (5%), *nvd*F*_ST_* is still able to detect the effect of selection in spite of the homogenizing effect of migration. The detection power ranges between 31-57% with a false positive rate of 0-0.1% (Table 5).

**Table 5.** Performance of *nvd*F*_ST_* in the short term (500 generations) with a single selective site. Selection was *α* = 600 and *Nm* = 50. Mean localization is given in distance (kb) from the real selective position.

| $\Sigma$ | $\theta$ | $\rho$ | %Power | %FPR ($\gamma = 5\%$) | $q$-value | Localization (kb) |
| --- | --- | --- | --- | --- | --- | --- |
| 116 | 60 | 0 | 47 | 0 | 0.0098 | $\pm 152$ |
| 180 | 60 | 4 | 57 | 0 | 0.0042 | $\pm 123$ |
| 178 | 60 | 60 | 31 | 0.1 | 0.0096 | $\pm 4$ |

$\Sigma$ : Mean number of shared SNPs per Mb. $\theta$ : Mutation rate. $\rho$ : Recombination rate. FPR: false positive rate. $q$ -value: mean estimated $q$ -value for the significant tests.

Concerning the EOS test, the power was below 10% with no false positives under this setting (see Discussion).

### HapMap Data

In order to check how the *nvd*F*_ST_* method works under real data, we have analyzed phased haplotypes from human chromosome 2. This chromosome has been widely analyzed by several methods, so we can also compare the performance of *nvd*F*_ST_* in this way. Concretely, we analyzed the chromosome 2 from northern and western European (CEU), East Asian (ASN: CHB + JPT), and Yoruba (YRI) human populations from Phase III of the International HapMap Project [21]. The data consisted in 116,430 SNPs in the unrelated samples downloaded from the project page (http://hapmap.ncbi.nlm.nih.gov/, last accessed May 29, 2016). The original data files, and the command line options for the analysis with HacDivSel, jointly with the obtained output files, are available in the S3_HapMap file.

The HacDiveSel program automatically filters the data, using only SNPs shared between each pair of populations. Therefore we analyzed 86,483 SNPs under the ASN-CEU comparison; 79,272 under the ASC-YRI, and 74,582 for the CEU-YRI. We set the program to select the 0.1% highest *nvd* values within each assayed window size. For these candidates, we only considered those having a positive value under the sign test (5), jointly with a significant *p*-value under the *F*_ST_ test.

In Table 6 (see also S3_HapMap file), we provide the genomic regions in which we found significant SNPs, and their corresponding genes within these regions. The estimated *q*-values were below 0.01 in every case. We found some of the highest *nvd* values, involving more than 40 SNPs, within the region 135-136 Mb in CEU-YRI comparison. This includes SNPs in a quite precise localization (135.78-135.83) of the lactase (LCT) gene jointly with some well-known linked candidates for recent selective sweeps as RAB3GAP1, ZRANB3, MCM6 and R3HDM1 genes [16, 17, 47]. The SNPs in this region showed high *F*_ST_ values (Table 6).

This result was expected as it is known that Yoruba does not show signature of selection at LCT while the signature is strong in CEU [16]. Recall however, that *nvd*F*_ST_* has no required any a priori information on candidate sites but just computed the *nvd* statistic for every SNP, performed the sign and *F*_ST_ tests and finally return those from the highest 0.1% *nvd* values that passed the sign test and had a significant *F*_ST_ value.

Another strong signal also occurred for the 218.9 - 219.4 region which includes genes previously reported under divergent selection as TTLL4 [17], as well as genes related to growth and immune maturation as NHEJ1 [48]. This region also presents some significant signal in the ASN-YRI comparison. Also within this region, and still under the CEU-YRI comparison, we detected the IL34 gene, which has been previously reported as giving strong signature of selection in the YRI population [49]. Another interesting signal in this region is at the SNP rs613539 which corresponds to SLC23A3 gene, which has been claimed to be a candidate for schizophrenia susceptibility in the Japanese population [50]. This SNP has been significant in CEU-YRI and ASN-YRI comparisons but not in the ASN-CEU one (Table 6 and S3_HapMap file).

Two more regions, including NBAS and CLASP1 genes, were detected under the CEU-YRI comparison. NBAS has also been detected in the ASN-CEU pair (see below). The CLASP1 gene is close to GLI2, already reported as a top region by the XP-CLR method [51]. In the ASN-YRI comparison, the region detected was 203.4-203.9, which is close to the NIF3L1 gene and includes a SNP previously reported under non-synonymous differentiation in these populations [17].

Finally, the significant *nvd* candidates with lowest *F*_ST_ values, correspond to the regions detected in the CEU-ASN comparison (this case also involves a lower number of regions). In this comparison, the highest *nvd* occurred in the NBAS region that had also the strongest signal in the CEU-YRI pair. Another interesting region is the 27.9-28.1 which includes the BRE gen (Table 6).

**Table 6.** Top significant divergent selection regions of the human chromosome 2 based on the *nvd*F*_ST_* test for the 0.1% highest *nvd* values.

| Populations | # significant SNPs | Positions (Mb) | $F_{ST}$ | Genes |
| --- | --- | --- | --- | --- |
| ASN-CEU | 67 | 15.49-15.54<br>27.95-28.1 | 0.12-0.17<br>0.21 | NBAS<br>RBKS, MRPL33, BRE |
|  |  | 63.6-63.98 | 0.15-0.21 | WDPCP, UGP2, ACA59 |
|  |  | 83.23-83.24 | 0.16 | Intergenic |
| ASN-YRI | 46 | 27.98-28.09 | 0.25 | MRPL33, BRE |
|  |  | 56.89 | 0.16 | intergenic |
|  |  | 135.69-135.72 | 0.23-0.35 | ZRANB3 |
|  |  | 203.4-203.9 | 0.18-0.29 | ICA1L, WDR12, CARF,<br>CYP20A1 |
|  |  | 117.28-117.30 | 0.22-0.36 | intergenic |
|  |  | 210.26 | 0.19 | MAP2 |
|  |  | 218.99-219.2 | 0.29-0.53 | CTDSP, VIL1, AK302678,<br>NHEJ1-XLF, SLC23A3, piR-39082, ABCB6, AK091345,<br>ATG9A |
| CEU-YRI | 116 | 15.52-15.53 | 0.29 | NBAS |
|  |  | 27.9 | 0.25 | RBKS-MRPL33 |
|  |  | 122.02-122.04 | 0.22-0.23 | CLASP1 |
|  |  | 135.4-136.9 | 0.29-0.63 | ACMSD, CCNT2-AS1,<br>MAP3K19, RAB3GAP1,<br>ZRANB3, LCT, MCM6,<br>R3HDM1 |
|  |  | 178.08 | 0.41 | AGP |
|  |  | 218.8-219.4 | 0.28-0.43 | ARPC2, LINC00608, VIL1,<br>USP37, NHEJ1-XLF,<br>RQCD1, SLC23A3, IL34,<br>ZNF142, BCS1L, GLB1L,<br>STK36, TTLL4, PTPRN,<br>CYP27A1 |

The positions correspond to the minimum and maximum locations for each given region. The $F_{ST}$ range corresponds to the minimum and maximum $F_{ST}$ for each region.

### *Littorina saxatilis* Data

To check the performance of the EOS test with real data, we provide an example with the aim of checking if even under its conservativeness is able of detecting some outliers on an already published dataset from the marine gastropod *L. saxatilis*. Recall that under the wide variety of scenarios assayed, EOS had a very low false positive rate. Thus the detected outliers, if any, may be good candidates, or at least good indicators, of the signal of divergent selection.

The rough periwinkle (*Littorina saxatilis*) is a marine gastropod mollusk that represents an interesting system for studying adaptive divergence and parallel speciation at different spatial scales. In Europe, *L. saxatilis* has adapted to different shore habitats resulting in both an exposed-to-wave, and other non-exposed (but crab-accessible) ecotype. The crab-accessible ecotype has large thick shells while the exposed-to-wave ecotype consists of smaller snails with thin shells and a larger shell aperture. Several experimental studies have shown that these ecotypes have been able of evolving local adaptation in the face of gene flow even at small spatial scales [52].

Ravinet *et al*. [53] have recently published a study where they used RAD loci as dominant markers to quantify shared genomic divergence amongst *L. saxatilis* ecotype pairs (wave vs crab) on three close islands on the Skagerrak coast of Sweden. These islands are connected by weak gene flow. In their outlier analysis, they filtered the data for sex-linked loci and null alleles. The filtering was applied for each island separately.

We applied the EOS test to analyze the separate-island filtered loci from Ravinet *et al*. Loci with null allele frequency equal or higher than 0.5 were discarded. We also discarded those polymorphisms not shared between ecotypes from the same location. Additionally, we required a minimum frequency allele of 4 per metapopulation sample size. Thus, we have excluded about 10-20% of the original individual-island filtered loci. The results of the between ecotypes outlier analysis using EOS are shown in Table 7. Results show that the number of outliers detected as significant in this study is much less than in the original study.

**Table 7.** Outliers detected after EOS analysis of the individual-island filtered loci from *L. saxatilis* data. Numbers in parentheses refer to the results in the original analysis [53].

| Island | Unique | Only with Jutholmen | Only with Ramsö | Only with Saltö | Shared all | Total |
| --- | --- | --- | --- | --- | --- | --- |
| Jutholmen | 7 (59) | — | 2 (13) | 0 (16) | 1 (9) | 10 (97) |
| Ramsö | 24 (86) | 2 (13) | — | 2 (21) | 1 (9) | 29 (129) |
| Saltö | 6 (134) | 0 (16) | 2 (21) | — | 1 (9) | 9 (180) |

We find a total of 48 outliers in the three islands while the number originally found was 406 [RAD loci in Table 2 of 53]. This is not surprising given the conservative nature and low false positive rate of EOS. However, note that we find a 2% (1/48) of SNPs shared by all islands which is quite similar to the 2.2% (9/406) found in the original study. Considering the islands by pairs, Jutholmen and Ramsö share 2 outliers, Saltö has no outlier in common with Jutholmen but it shared 2 with Ramsö.

For the outliers detected by EOS, the mean *F*_ST_ between ecotypes ranges between 0.48-0.6 (Table 8). The *q*-values are high (0.51 - 0.76) although we already know by the simulations that this may indicate linkage between the markers, more than a real false positive rate [see also 11].

**Table 8.** Summary of EOS analysis for the between ecotypes *L. saxatilis* data [53].

| Island | Non-outliers | Outliers not in EOS | EOS | $F_{ST}$ | $F_{ST\_EOS}$ | $pval_{EOS}$ | $qval_{EOS}$ |
| --- | --- | --- | --- | --- | --- | --- | --- |
| Jutholmen | 4564 | 112 | 10 | 0.045 | 0.48 | 0.002 | 0.51 |
| Ramsö | 4602 | 82 | 29 | 0.064 | 0.53 | 0.005 | 0.63 |
| Saltö | 4632 | 51 | 9 | 0.060 | 0.60 | 0.002 | 0.76 |

$F_{ST}$ : Mean $F_{ST}$ for the analyzed loci. $F_{ST\_EOS}$ : Mean $F_{ST}$ for the loci included in the extreme outlier set. $pval_{EOS}$ : Mean $p$ -values across the loci included in the extreme outlier set. $qval_{EOS}$ : Mean $q$ -values across the loci included in the extreme outlier set.

## Discussion

The aim of this study was to develop two complementary strategies, haplotype-based and outlier-based, for the detection of divergent selection in pairs of populations connected by migration. Because high rate of false positives is a known concern of outlier-based methods [9-11], the proposed methodology was especially designed to minimize the false positive rate. Additionally, both methods should be useful for non-model species and therefore, it should not be necessary either to have a priori functional information on candidate regions, or to perform neutral simulations to obtain critical cut-off values.

We have shown that *nvd*F*_ST_*, which combines haplotype-based and *F*ST-differentiation information, is a powerful strategy for detecting divergent selection. The method has proven to work well even when the phasing accuracy is not perfect. However, the *nvd*F*_ST_* algorithm does not perform well when the whole set of markers segregates independently. To deal with the latter, a second method was proposed based on the idea that the outliers caused by the effect of divergent selection would cluster apart from those caused by different demography issues. This extreme outlier set test EOS, was intended to be conservative because of the aforementioned tendency of outlier-based methods to produce false positives. Under the simulated scenarios, the EOS test behaves well when markers are independent or under weak linkage, reaching powers between 60-90% while maintaining false positive rates below the nominal level.

### Polygenic Architecture

In general, the *F*ST-based methods cannot detect selection in polygenic scenarios [8, 11] because these tests are specifically designed for finding *F*_ST_ values larger than average, which are difficult to discover if the frequency differences are slight for the polygenic loci, or if the overall *F*_ST_ is high. On the contrary, the *nvd*F*_ST_* performs even better under such scenario. The explanation for this good performance is that the distributed selective signal facilitates the discovery of the corresponding haplotype patterns of divergent selection. These patterns are coupled with the occurrence of high frequency at the target site in one population and low in another. The good performance of *nvd*F*_ST_* under the polygenic setting, therefore, occurs because a) we select concrete SNPs because of their selective pattern and b) this coincides with higher *F*_ST_ at these SNPs.

### Position Effect, Positive Predictive Value and FDR by site

Besides the detection of the signal of selection, we have also inferred the location of the selective site. It has been shown that under *nvd*F*_ST_*, the localization is better when the selective site is at the center of the chromosome. The ability of localizing the selective position is still a pending issue for many of the selection detection methods. There is also plenty of room for improvement under the *nvd*F*_ST_* and EOS methods in this regard, for instance by exploring the relationship between recombination and the window sizes that yield the highest scores. Indeed, the interplay among divergent selection, recombination, drift and migration should be considered for further improving the efficiency of the methods.

Concerning true and false positive rate measures, we have given a) the true positive rate (power) in terms of the percentage of runs in which selection (at any site) was detected from simulated selective scenarios and b) the false positive rate as the percentage of runs in which selection (at any site) was detected from simulated neutral scenarios.

The error rate can also be measured by the number of sites falsely detected as selective. First, it should be clarified that under the neutral scenario, we always considered a false positive when any site within the chromosome has been signaled as selective. Thus, when we are saying that FPR is below 5%, in more than 95% of the neutral files no single site was detected as selective, and so the error rate by site is 0. Therefore, we can consider the number of sites advocated as selective in those few runs in which neutrality was falsely rejected. In the case of the EOS test, this number was just 1 single site per Mb when markers are independent or 1 to 4 when they are linked.

In the case of *nvd*F*_ST_*, the situation is different just because various window sizes are assayed and a number *k* of candidates is considered, so that the maximum number of false positives by genome can be at most *w* × *k*, where *w* is the number of different window sizes. However, the number of false positives would also be dependent on the recombination rate.

In most cases, we have assayed 2-4 window sizes and just 1 candidate, and the average number of falsely detected sites in the neutral files was close to 1.

When a higher number of candidates (*k*>1) was studied the percentage of false positives was always less than γ × *k* (γ = 0.05). For example, in the extreme case of inspecting the 100% of neutral sites from an ultra-high density linkage map (about 60,000 candidates with ρ = 120), we found 1-2% false positives independently of the window size. For the selective scenarios, we are also interested in the per site comparison, i.e. the proportion of detected sites that lies at, or close to, the true selective position. The linkage relationship will depend on the recombination rate and so we would measure the distance in centimorgans (cM). Therefore, if we consider a given position as being true only when it lies at distance of less than *t* cM from the real position, then the positive predictive value (PPV) can be measured as PPV = #(detected sites at distance ≤ *t*) / # detected sites. Similarly, the false discovery rate per site is, FDRl = 1 -PPV.

Thus, given the positions obtained in our simulations, when we set *t* = 0.15 cM the PPV is in most of the cases 100%, i.e. all detected positions lie within the acceptance region, except for some cases under the highest recombination rate (ρ = 60). In the latter cases under the *nvd*F*_ST_* test, the PPV decreases to 85-95% with a worst case of 70% when the selective site is located at the very extreme of the chromosome. Therefore, in these latter cases (*nvd*F*_ST_*; ρ = 60), when claiming that selection has been detected, we are assuming that 5-15% of the times the detected position could be far away from 0.15 cM. However, If we assume as correct any position within 1 cM distance from the true one (*t* = 1), then the per-site FDR_l_ is virtually zero in every studied case. The latter means that any detected site lies closer than 1 map unit from the real position.

### High Migration Scenario

It is important to note that under high migration, *nvd*F*_ST_* maintains a reasonable power. However, the power diminishes with the highest recombination rate. This may occur due to the combined effect of gene flow and recombination, that generates intermediate HAC mean values *m*_1_ and *m*_2_ and similar variances. Indeed, for a given selection intensity, the higher the *Nm* requires tighter linkage for the establishment of divergent alleles [54].

In the case of the EOS test, there is an obvious tradeoff between the stringency of the cutoff point for the outlier set and the migration rate. The cutoff depends on the *F*_ST_ upper-bound which is a function of the number of populations, the sample size and the minimum allele frequency. However, this does not take into account the effect of migration. A possible solution to improve the efficiency of EOS under high migration would be to update the cutoff point as a function of the migration rate.

### HapMap data

The applicability of the *nvd*F*_ST_* method to real data was evaluated by analyzing the human chromosome 2. For each pair of populations, the full set of SNPs in the phased haplotypes was directly analyzed without any assumption about specific candidate regions or reference SNPs.

Our method successfully detected the lactase persistence region that is known to expand over more than 1 Mb in chromosome 2 of European populations [47, 55]. In addition, we detected other regions previously reported as under ongoing selective sweeps in human populations (see Table 6 and S3_HapMap file), together with another that has not been explicitly reported from the HapMap comparisons. Worth mentioning is the SLC23A3 gene that has been associated to schizophrenia susceptibility in the Japanese population [50] and that we found to be significant both in ASN-YRI and CEU-YRI comparisons.

We also found as selected other regions not previously reported. For example, in the ASN-CEU comparison, 20 SNPs had high and significant *nvd* value within the NBAS gene. Mutations in this gene has been associated to short stature and different multisystem disorders [56, 57]. Again in the ASN-CEU comparison, the BRE gene showed quite high *nvd* values. This gene encodes a component of the BRCA1-A complex. These results illustrate how the *nvd*F*_ST_* method can be used in exploratory studies to detect locally adapted polymorphisms that could be interesting candidates for association studies.

### *Littorina saxatilis* Data

The natural systems where local adaptation occurs can be of great complexity [58]. Local adaptation may occur most likely due to alleles with large effect but also under a polygenic architecture [58, 59]. In addition, the geographic structure and the migration-selection balance can generate complex patterns of the distribution of genetic variation [60]. The *L. saxatilis* ecotypes are an especially interesting system to study local adaptation in presence of gene flow [61]. This system has an exceptional level of replication at different extents, such as country, localities within country, and the micro-geographical level of the ecotypes. In the case of the Swedish populations, the pattern of differentiation can be separated in factors such as, localities and habitat variation among islands —that may be caused by genetic drift— and variation between habitats within localities –that may be caused by divergent selection [61]. There are also different mechanisms by which parallel adaptation may occur, resulting in different predictions about the proportion of shared adaptive variation among localities.

Regarding the shared genomic divergence of the *L. saxatilis* system in Swedish populations, it seems to be a small proportion of the total genomic divergence [53, 61, 62]. That is, the majority of the genomic variation linked to the evolution of ecotypes is not shared between the studied islands. The EOS analysis of the Ravinet *et al*. data seems to support this finding. At the same time, we identify far fewer outliers, being Saltö —which is closer to the mainland— the island with the lowest number. However, this result is the opposite of the result in Ravinet’s study, where Saltö had the highest number of outliers. This difference can be caused by an excess of false positives in the original study, though we cannot rule out that our findings can be an artefact due to the conservativeness of EOS.

As a final remark, the strategy used in *nvd*F*_ST_* is in line with the suggestion of combining multiple signals from different tests as a way of improving the power/false positive rate relationships for the selection detection methods [18, 63, 64]. Accordingly, the *nvd*F*_ST_* test does this. It combines haplotype and population differentiation information and may be a helpful tool to explore patterns of divergent selection when approximate knowledge of the haplotype phase is available. Alternatively, the EOS method is a conservative outlier test useful when the full set of SNPs is unlinked or under weak linkage. Both strategies can be applied without the need of performing neutral simulations and have a low false positive rate.

## Acknowledgements

I would like to thank E. Rolán-Alvarez, C. Canchaya and Auriel Sumner-Hempel for useful comments on the manuscript.

## Supporting Information

S1_Appendix. Format: PDF. Description: Supplementary Appendix (from A1 to A7 includes Figures S1-S3 and Tables S1 and S2).

S2_Sim. Format: rar. Description: Input files for the GenomePop simulations and the command lines for their analysis with HacDivSel.

S3_HapMap. Format: rar. Description: the original data files from HapMap as well the command line options for the analysis with HacDivSel, jointly with the obtained output files.

S4_HacDivSel_program_manual. Format: PDF. Description: Complete manual for the program that implements the methods described in the paper.

